# Bumblebees locate goals in 3D with absolute height estimation from ventral optic flow

**DOI:** 10.1101/2024.09.02.610798

**Authors:** Annkathrin Sonntag, Martin Egelhaaf, Olivier J. N. Bertrand, Mathieu Lihoreau

## Abstract

When foraging, flying animals like bees are often required to change their flight altitude from close to the ground to above the height of the vegetation to reach their nest or a food source. While the mechanisms of navigating towards a goal in two dimensions are well investigated, the explicit use of height as a source for navigation in three dimensions remains mostly unknown. Our study aims to unravel which strategies bumble-bees use for height estimation and whether they rely on global or local cues. We expanded a 2D goal localization paradigm, where a goal location is indicated by cylindrical landmarks, to the third dimension by using spherical landmarks to indicate a feeder’s position in 3D and examined the search pattern of bumblebees. Additionally, we assessed the ability of bees to estimate the height of a feeder based on local landmarks and global references such as the ground floor. The search distribution for a feeder’s position in 3D was less spatially concentrated compared to in 2D. Assessing the bees’ height estimation ability, we found that bees could estimate a feeder’s height using the ground floor as a reference. However, the feeder needed to be sufficiently close to the ground floor for the bees to choose correctly. When bumblebees are faced with the challenge of foraging in a 3D environment where the height of a food source and landmark cues are important, they demonstrate the ability to learn and return to a specific flower height. This suggests they rely on ventral optic flow for goal height estimation in bumblebees.

## INTRODUCTION

Bees have remarkable navigational abilities in three-dimensional space (Brebner et al., 2021; Bullinger et al., 2023; Menzel, 2023; Osborne et al., 2013; Woodgate et al., 2016). While previous research has extensively explored cue utilization and navigation strategies in two dimensions (Buehlmann et al., 2020; Collett et al., 2013; Kheradmand et al., 2019), bees fly at various heights. For example, they feed on flowers ranging from close to the ground to blossoms on trees at the height of multiple meters. They can also fly at heights of hundreds of meters over more considerable distances (Dillon et al., 2014; Gibo, 1981). This provides the bees with extra visual cues associated with altitude that they could use for accurate navigation.

Height estimation involves two key components: determining one’s own altitude and estimating the height of a goal. Insects use multiple strategies for altitude estimation, including optic flow—the rate of image motion across the retina, which depends on distance and velocity (Srinivasan et al., 1991; Esch et al., 1996). However, relying solely on ground-based optic flow can be unreliable, as changes in speed alter perception (Srinivasan et al., 1996; Baird et al., 2005). To compensate, insects may use additional cues such as texture gradients or optic flow from nearby objects (Linander et al., 2016).

Mechanisms for height estimation are crucial not only for determining the insect’s own flight height but also for estimating the height of goals, such as food sources like blossoms of trees or the home like the nests that can be located above the ground in trees. In principle, estimating the height of goals may be accomplished in different manners. The insect could use absolute height relative to the ground (i.e. a ground-based allocentric estimation of the height of a goal) or the height of the goal relative to their flight altitude (i.e. an egocentric estimation of the goal’s height). An allocentric estimation may not only be ground-based, but also relative to objects. In such a case the vertical distance between objects and the goal would be estimated (i.e. an object-based allocentric estimation). Studies have demonstrated that honeybees can accurately estimate height on a small spatial scale of a few centimeters (Lehrer et al., 1988; Srinivasan et al., 1989). During a transfer test, bees trained on the highest flower at 5 cm chose the flower at the trained height of 5 cm instead of the highest flower in the test at 10 cm. Thus they may have used a ground-based allocentric estimation or an egocentric estimation of the goal, but do not seem to use an object-based allocentric estimation at this scale. This ability of bees to discriminate height at a small spatial scale raised the question on whether and how they use such information to reach a 3D goal at a larger scale.

Studies in 2D indicate bees can learn the position of a goal using landmarks. Honeybees were trained to learn a feeder position on the floor surrounded by three cylinders (Cartwright et al., 1983; Cheng et al., 1987). When the feeder was removed, the bees showed a concentrated search at the previous location of the feeder using landmarks as reference points. Movements and cue utilization in 2D and 3D environments pose distinct challenges to navigating organisms. While navigational strategies may be similar on a 2D plane, navigating in 3D introduces additional complexities due to the need to account for vertical movement and more significant potential for error possibilities. For instance, a study comparing bees’ preference for flowers arranged horizontally versus vertically revealed differing performance levels based on cue presentation orientation (Wolf et al., 2015). This suggests that bees exhibit variations in performance when searching for food based on cues presented in different spatial orientations.

In this study, we extended the 2D landmark-based navigation paradigm (Cartwright et al., 1983; Cheng et al., 1987) into three dimensions. We asked whether bumblebees can learn a feeder position in 3D and whether they estimate the goal height based on absolute distance to ground vs relative distance to landmarks. We hypothesized that bees trained to locate a feeder using landmarks would search around the same 3D location after the feeder was removed. moved. As the results of the first experiment deviated from this expectation, we hypothesized that the bees may use an absolute height estimate of the feeder relative to the ground and use relative height information of the spherical landmarks relative to the feeder. By analyzing the flight trajectories and search behavior of bumblebees, we provide new insights into goal localization while focusing on height estimation of flying insects in 3D environments.

## METHODS

### Animal handling

Three *Bombus terrestris* colonies were used, provided by Koppert B.V., France. We tested one colony after the other from September 2022 to January 2023. The bees, arriving in a small box, were transferred under red light (not visible to the bees (Skorupski et al., 2010) into a dark gray acrylic box (0.24 × 0.24 × 0.4 m) with a transparent lid easing the monitoring of the colony health. To simulate natural underground lighting conditions (Goulson, 2010), the hive was covered with a black cloth.

We provided pollen balls *ad libitum* in the hive box. For these pollen balls, 50 ml commercial ground pollen collected by honeybees (W. Seip, Germany) were mixed with 10 ml water. Sugar water, a sweet aqueous solution (30% saccharose, 70% water in volume), was provided *ad libitum* to the bees in a micro-gravity feeder in the flight arena. The micro-gravity feeder consisted of a falcon tube screwed on a 3D-printed blue landing platform with small slits where the bees could land on and suck the sugar solution out of the small slits. The landing platform of the feeder had a diameter of 6 cm. The bees were tagged with individually numbered plastic tags (W. Seip, Germany) glued on their thorax with melted resin for individual identification.

### Flight arena

The flight arena was a 4 m x 4 m x 2 m windowless indoor room. The floor was covered with a red and white pattern with a 1/f frequency pattern (pink noise), mimicking the natural surroundings (a distribution observed in nature; (Schwegmann et al., 2014)), providing enough contrast for the bees to use optic flow. The walls and the ceiling were covered by white tarpaulin (for photographs see Supplementary Material). The light was provided by 18 light bulbs placed behind the white tarp. These lighting conditions were sufficient for the bees to fly naturally, as indicated by their recorded 3D trajectories (see 1C and 6). The colony was connected via small boxes (8 × 8 × 8 cm) to a flight arena. The small boxes had closable doors to select bees individually. The bees could access the flight arena via a small tube (2.5 cm diameter) in one corner (Fig. 2A) at a height of 1.25 m. The experimenter could access the arena through a cutout door in the tarpaulin fixated with velcro. Once we saw regular traffic of bumblebee foragers between the colony and the feeder, the experimental tests were started. For the tests, the bees were manually removed from the flight arena and only one bee at a time was allowed to enter it using doors at the colony entrance tube. One bee at a time was allowed into the arena and its search for the feeder was recorded for three minutes. After entering the arena, the bee walked on a platform at a height of 1 m and had to take off that platform to enter the flight arena. The bee had a maximum of two minutes to take off otherwise the trial was discarded. Each bee was tested only once per recording session, with one session in the morning and another in the afternoon.

### Video tracking

The bees’ flight trajectories were recorded inside the flight arena using four synchronized Basler acA 2040um-NIR cameras with a frequency of 62.5 Hz (as in Sonntag et al., 2024). The cameras were positioned in the corners of the arena, facing upwards toward its center. Recordings began before a bee entered the arena, with the first 60 seconds used to calculate a background image (bee-free). Subsequently, only cropped sections of the frames (42 × 42 pixels) showing significant differences from the background were saved, along with their positions in the image. The recording script was written in C++.

These image crops were analyzed using a custom neural network written in python (3.8.17) to classify whether they contained a bee. The network was a five-layer feed-forward architecture with three convolutional layers (40, 64, and 32 filters), followed by a dense layer and a single-neuron output layer encoding the probability of the crop containing a bee. Rectified Linear Unit (ReLU) activation functions were used throughout. The model, trained on approximately 1 million manually classified images over 10 epochs, achieved nearly 100% accuracy. The trained model, along with the database and complete pipeline—from labeling to training, evaluation, and application—is publicly available (Bertrand et al., 2024b; Bertrand et al., 2024a).

To reconstruct 3D trajectories, the cameras were calibrated to determine their intrinsic and extrinsic parameters. We aligned the cameras to a shared coordinate system. The cameras were synchronized to capture frames with the same time stamps. We combined 2D positions from at least two cameras using triangulation. Only triangulated positions with a reprojection error below 10 pixels were used. The median position across cameras was calculated for each frame. The reprojection error is the distance between the originally measured position and the reconstructed 3D position reprojected back onto the original image. The recording and triangulation pipelines are publicly available (Bertrand et al., 2024c).

Finally, the 3D trajectories were reviewed for quality. Instances of non-biological speeds (above 4 m/s Goulson, 2010) or positions outside the arena were flagged, and neighboring crops were manually inspected. Data analysis was performed using Python (3.8.17).

### 3D goal localization

We adapted a 2D goal localization paradigm to investigate how bumblebees locate a 3D goal position indicated by landmarks (Cartwright et al., 1983; Cheng et al., 1987; Collett et al., 2013; Doussot et al., 2020; Wehner et al., 1996) (Fig. 1). The bees (N = 13) were trained to a feeder position surrounded by three landmarks. Then, we tested if they could find the feeder position based on the landmark cues after the feeder was removed (Fig. 1A&B). We hypothesized that after learning a feeder position indicated by landmarks, the bees would search in a volume concentrated around the previous position after the feeder was removed. Three acrylic spheres (diameter of 8 cm) were placed in one corner of the arena surrounding a micro-gravity feeder (a plate with small slits and a 50 ml Falcon tube containing the sugar water). The landing platform of the feeder was placed at a height of 1.20 m above ground.

**Fig. 1.**
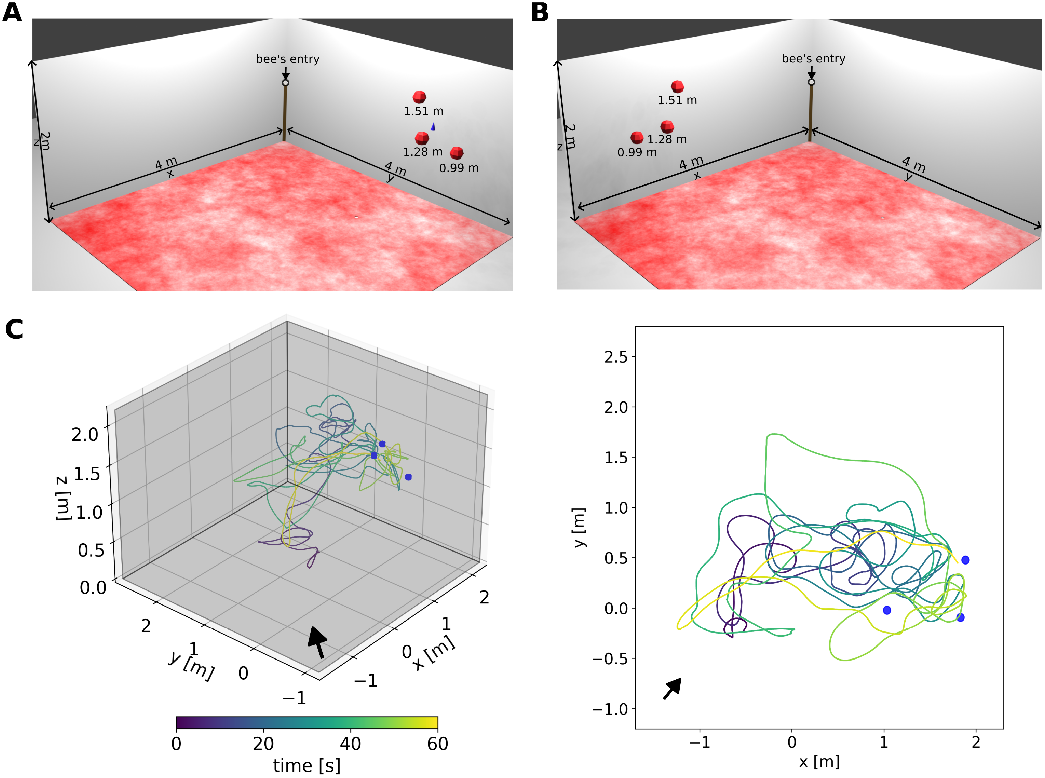
Flight arena with the landmark constellations for 3D goal localization experiment. **A** Schematics of the 3D goal localization setup for the training with red spheres as landmarks around the feeder, indicated by the blue cone. In each corner of the arena, one camera was positioned. The bees entered the arena through a tube at the top of the wooden stick in one corner (brown bar). The floor was covered by a red and white pattern (perceived as black and white by the bees (Skorupski et al., 2010)) to provide enough contrast for the bees to use optic flow for flight control. **B** Schematics of the setup for the shifted constellation test, similar to the training setup but the landmark constellation was shifted to the other side of the room and the feeder was removed. **C** An example flight in the shifted constellation test. The landing platforms are indicated in blue and the arrow shows the bees’ entry to the flight arena. The trajectories are colour coded by the time, blue indicating the entry to the area and yellow after 60 seconds of flight.

**Fig. 2.**
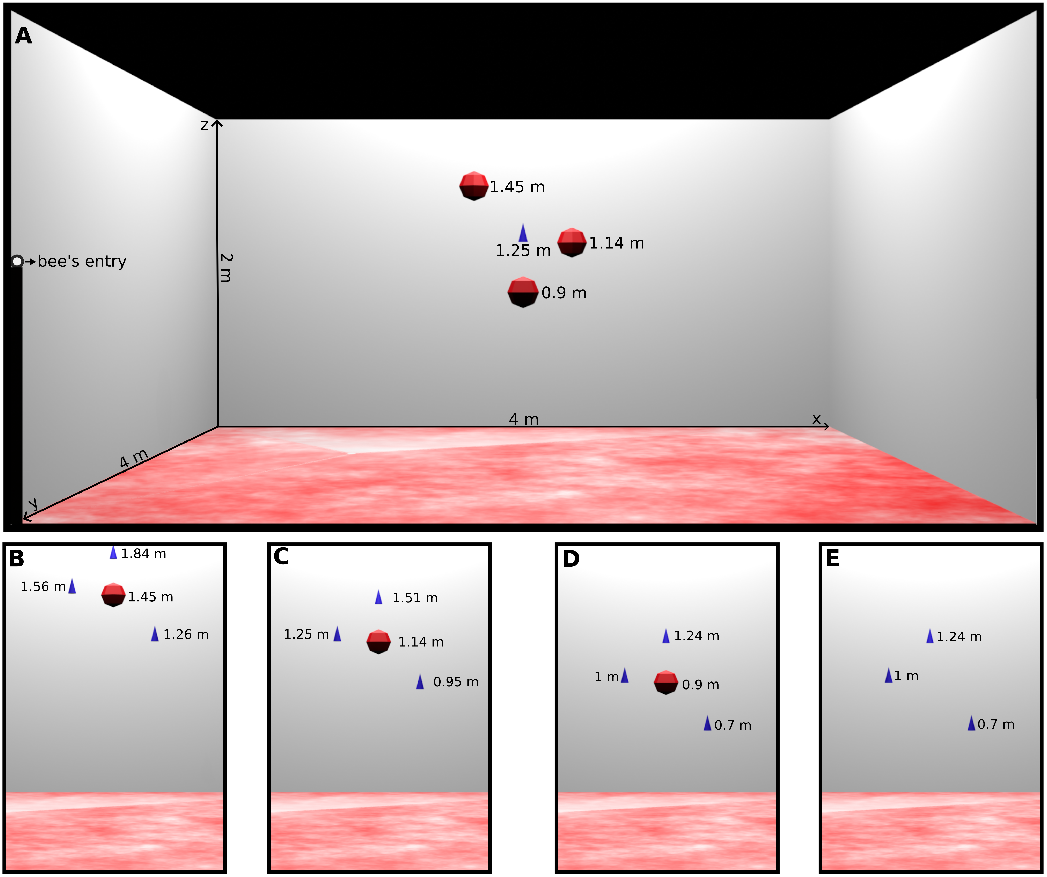
Schematics of the setup for the height estimation experiment with the training and test conditions. **A**: Flight arena for the height estimation experiment with the training constellation. **B-E**: The three test constellations with only one of the spherical landmarks (indicated by red circles) and three landing platforms (indicated by blue cones). Test conditions: high sphere (**C**), intermediate sphere (**D**),low sphere (**E**) and feeders-only (**F**).

The distance between the feeder and each of the spheres was similar (0.34 m, 0.37 m, 0.35 m) and they were hanging at three different heights (1.51 m, 1.28 m and 0.99 m). The spheres were coloured in red to be perceived as black by the bees (Skorupski et al., 2010) but provided enough contrast for tracking the bee in front of them with the cameras. These dark-colored objects (dark blue for the feeder and red, perceived as black, for the spheres) provided strong contrast against the white background of the walls, making them clearly perceivable for the bees (Giurfa et al., 1996; Meyer-Rochow, 2019).

Given that bumblebees can detect objects within a visual angle of approximately 2.3–2.7 degrees (Dyer et al., 2008), the spheres were visible from most positions within the arena, except when the bees were near the walls. In contrast, the feeder was visible only from over a meter away (see Supplementary Material). This suggests that bees may first use the spheres as cues before locating the precise feeder’s position and height. Moreover, since we analyzed the bees’ search behavior over a period of two minutes, they had sufficient time to explore the arena and approach the cue constellation within a perceivable range.

During the training condition the bees could travel freely between the feeder and the colony and no recordings were made (Fig. 1A). During the test, we shifted the landmark constellation to see if the landmarks and not any other cues there were unavoidable in the arena were used to localize the feeder (Fig. 1B). We calculated the time the bees spent in the area of the training and shifted constellation (spherical areas with a radius of 0.53 m around the feeder, including the center of the spheres). We used the Mann-Whitney U test to compare if the bees spent more time at the shifted constellation than at the training position. This finding is consistent for radii between 0.1 to 0.6m around the feeder (see Supplementary Material).

### Height estimation

In the 3D goal localization experiment, the bees did not exhibit the expected concentrated search at the feeder location. To address the potential reason for this finding, we modified the setup and ran a second experiment. We introduced multiple feeders (landing platforms without any reward) at different heights during the test phase. This adjustment aimed to clarify whether bees rely on absolute height estimation relative to the ground or relative height information provided by spherical landmarks.

During the training phase, bees were presented with a single feeder surrounded by three spherical landmarks, all positioned equidistantly from the feeder in the horizontal plane (Fig. 2A). The landmarks were arranged at different vertical heights: one above the feeder (1.45 m, high sphere), one at the same height as the feeder (1.25 m, intermediate sphere), and one below the feeder (0.9 m, low sphere). By training bees with this configuration, we ensured they experienced a consistent spatial relationship between the feeder and the spheres, allowing them to use either absolute or relative height cues during the test phase. The feeder and spheres were centered in the arena to minimize potential bias from global environmental cues, which we attempted to remove as thoroughly as possible.

In the test phase, the design was altered: instead of one feeder and three spheres, bees were presented with three landing platforms as alternative landing sites (unrewarded feeders) and either one sphere or none (Fig. 2 B-E). The landing platforms were placed at the same planar (xy) locations as the training-phase spheres but varied in height. This allowed us to test whether bees could distinguish the landing platforms based on height alone, independent of their planar positions.

Including only one sphere in some tests was crucial to determine whether bees used relative height estimation between the sphere and the ground as a reference for locating the feeder. By placing the sphere at the center of the arena (at the planar location of the training feeder), the bees could not use cues based on their horizontal (xy) position in the arena to identify the feeder height. Instead, they had to rely solely on vertical height information. We also tested a condition without any sphere (“feeders-only” test) to see if bees could use absolute height cues without a local height reference.

The following four conditions were tested, each differing in the height of the sphere (if present) and the placement of the feeders:

- High-sphere test: The highest sphere (1.45 m) was presented, with feeders at 1.84 m, 1.56 m, and 1.26 m (Fig. 2B).
- Intermediate-sphere test: The intermediate sphere (1.25 m) was presented, with feeders at 1.51 m, 1.25 m, and 0.95 m (Fig. 2C).
- Low-sphere test: The lowest sphere (0.9 m) was presented, with feeders at 1.24 m, 1 m, and 0.7 m (Fig. 2D).
- Feeders-only test: No sphere was presented, but the feeders were placed as in the low-sphere test (Fig. 2E).

To eliminate potential biases from chemical markings, the landing platforms were cleaned with 70% ethanol before each recording (Cederberg, 1977; Foster et al., 1989; Chittka et al., 1999; Eckel et al., 2023). To assess whether the bees were searching for the feeder, we analyzed their speed and sinuosity in the regions around and between the feeders using the Mann-Whitney U test. A lower speed and higher sinuosity would indicate search behavior. We then compared the search time across feeder positions for each condition using a one-way ANOVA with Tukey post-hoc tests.

## RESULTS

### 3D goal localization

We hypothesized that bees can learn the position of a feeder in 3D relative to surrounding landmarks. To test this hypothesis, we modified a 2D goal localization paradigm. We replaced the cylindrical landmarks with spherical ones and hung the feeder (i.e. the goal) between the landmark constellation. For the control test, the spherical landmarks remained at the same position as during training, but the feeder was removed. We investigated whether bees show a concentrated search at the 3D position of the feeder relative to the landmarks in 3D, even if the feeder itself was removed. The bees searched preferentially in the area close to the landmark constellation and not anywhere in the arena (Fig. 3). However, contrary to our expectations, the kernel density probabilities of the bees’ search showed that the bees spent much time just before the low sphere which was also placed closest to the bees’ nest entrance (Fig. 3). The bees did not search in the center of the constellation where the feeder was placed in the training situation, but rather undershot the position by flying not far enough (Fig. 3**A**). The kernel density probability showed a peak just below the highest sphere on the z-axis (Fig. 3**B**&**C**).

**Fig. 3.**
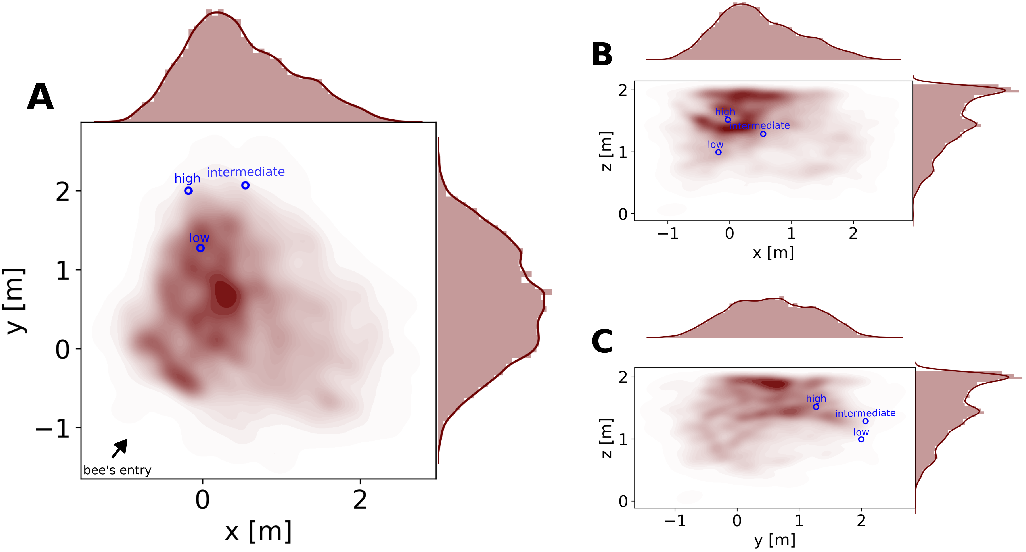
Search probability for the control test. The three subfigures show the kernel density probability (given by the color red, the darker the color is the more often the bees visited the position) of the bees’ flights in the arena for the control test. The positions of the spheres are indicated by blue circles. **A** shows the bees’ flight on the xy plane, **B** on the xz plane and **C** on the yz plane. At the top and right margins, histograms and probability density functions along the axes are shown.

For the test with the shifted constellation, the spheres were moved to the other side of the arena. This minimized any unintended cues in the arena. We investigated whether bees show a concentrated search at a 3D position of a feeder relative to the landmarks in 3D, even if the feeder itself was removed (Fig. 4). An examplary flight trajectory is shown in Figure 1D. The bees searched around the spheres, not in the empty corner where the constellation was placed during training (Fig. 5, Mann-Whitney *U*-test, *n*1 = *n*2 = 13, *p* = 0.0005, cohen’s *d* = 1.498). As in the control test, we observed some exploratory behavior in the center of the arena, just below the ceiling. This might have been caused by the artificial lighting above the ceiling (Fig. 4).

**Fig. 4.**
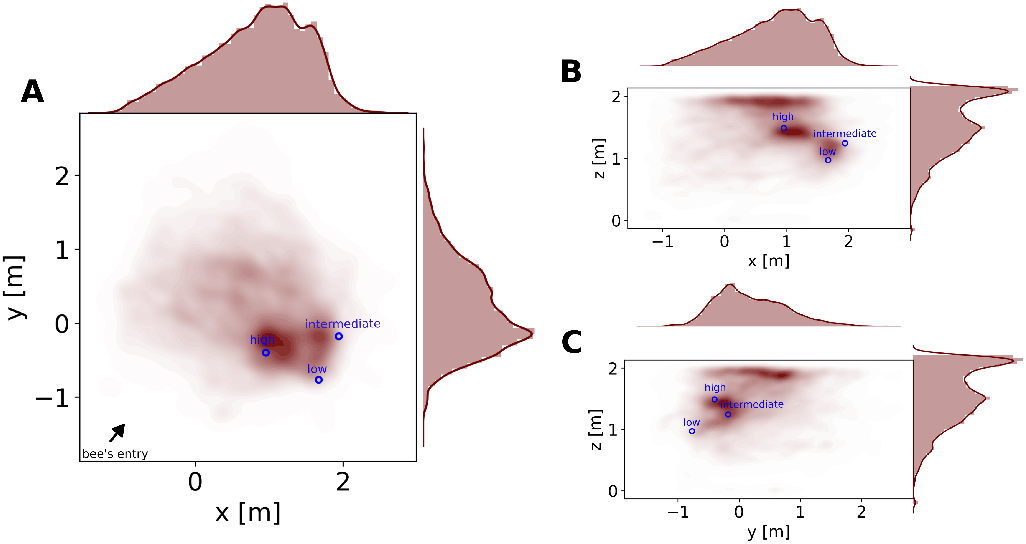
Search probability for the test with the shifted constellation test. The three subfigures show the kernel density probability (given by the color red; the darker the color is, the more often the bees visited the position) of the bees’ flights in the arena for the test with the control test (all three spheres shifted to another position). Blue circles indicate the positions of the spheres. **A** shows the bees’ flight on the xy plane, **B** on the xz plane and **C** on the yz plane. At the top and right margins, histograms and probability density functions along the axes are shown. **E** The search percentage of the bees at the two locations of either the training or the shifted test position of the landmark constellation.

**Fig. 5.**
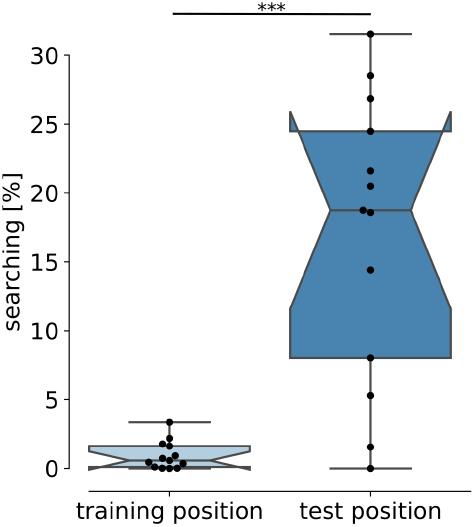
Comparison of the time the bees spent at the training and test position of the sphere constellation. The time spent at the training and test position for each bee is shown as black circles. The hatched boxplots display the median and the whiskers showvthe lower and upper range of 1.5 times the interquartiles. The bees searched more at the position of the shifted test constellation than at the training position (Mann-Whitney *U*-test, *n*_1_ = *n*_2_ = 13, p *<* 0.001).

### Height estimation

The 3D goal localization experiment showed that the bees search around the landmark constellation, but their search was not very concentrated around the goal. We changed our experimental paradigm to determine whether the bees could not localize the goal accurately based on landmarks or could not land at the goal, which differs from experiments with a ground-based goal. We therefore also offered the bees landing platforms in the test. Not just one as in the training, but three alternatives, which were in different spatial positions in relation to the landmarks (Fig. 2**A-D**). These landing platforms could be only differentiated by their height. We hypothesized that the bees would select the landing platform corresponding to the training situation by using the relative height between the sphere and the floor.

First, we tested if the bees were searching at the landing platforms. Bees lower their speed and fly in loops or sinuous paths when searching (Lihoreau et al., 2016). We tested their flight speed and sinuosity at the feeders and between them. We found a lower speed (Mann-Whitney *U*-test, *n*_1_ = *n*_2_ = 13, p = 1.663*e*^−38^, cohen’s *d* = -1.302, Fig. 7A) and higher sinuosity (Mann-Whitney *U*-test, *n*_1_ = *n*_2_ = 13, p = 2.352*e*^−08^, cohen’s *d* = 0.403, Fig. 7B) in the areas around the feeders. A more tortuous path indicates search behavior, while a straighter path suggests goal-directed behavior. Thus, we can conclude that the bees were searching at the feeders and quickly transitioning between them. Therefore, comparing the time spent at each feeder will be used as a quantitative indicator if they can discriminate the training feeder height from other heights.

During the training phase, a single feeder was paired with three spherical landmarks positioned at different heights: one higher, one at a similar height, and one lower relative to the feeder. This setup aimed to allow bees to associate the feeder’s location with the relative heights of the surrounding landmarks.

To investigate whether the bees learned the relationship between the feeder and individual spheres, we conducted three tests, each using only one of the spherical landmarks: high-sphere, intermediate-sphere, or low-sphere. In these tests, we provided three landing platforms as alternatives (unrewarded feeders) at different heights. These heights corresponded to the positions of the feeder during training—higher, similar, or lower relative to the height of the sphere used in each test.

We hypothesized that if the bees had learned the feeder’s position relative to the spheres, they would search for the feeder at specific heights corresponding to each sphere’s relative position. For example, with the high sphere, they would search at the lowest platform; with the intermediate sphere, at the middle platform; and with the low sphere, at the highest platform. Exemplary flight trajectories for each test condition are shown in Figure 6.

**Fig. 6.**
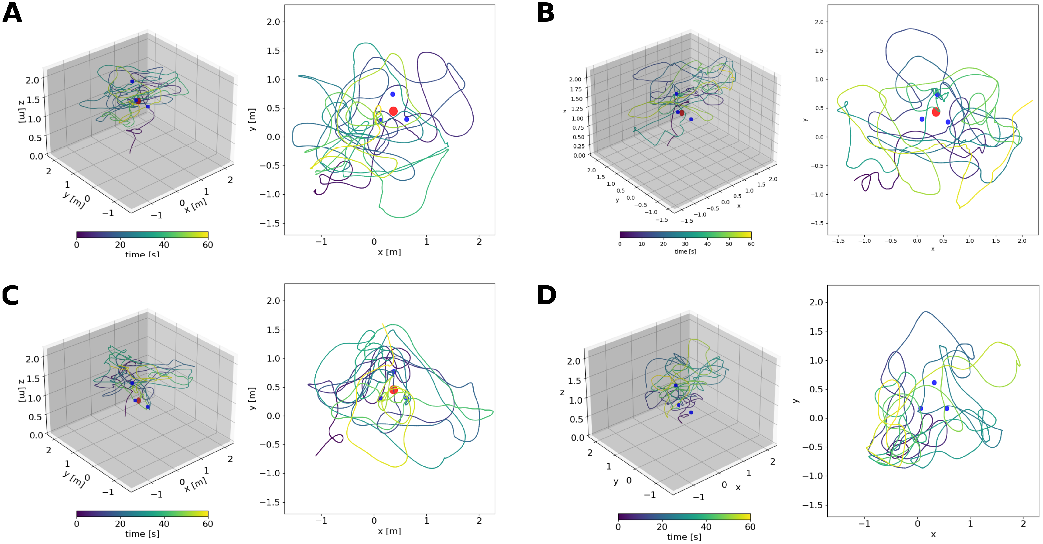
An examplary trajectory for each test of the height estimation experiment (high sphere (**A**), intermediate sphere (**B**), low sphere (**C**) and the test with only feeders without the low sphere (**D**). The left panel of each subplot shows the 3D trajectory with the red sphere and blue feeders. The trajectories are colour coded by the time, blue indicating the entry to the area and yellow after 60 seconds of flight. The bee’s entrance is indicated by a black arrow.

**Fig. 7.**
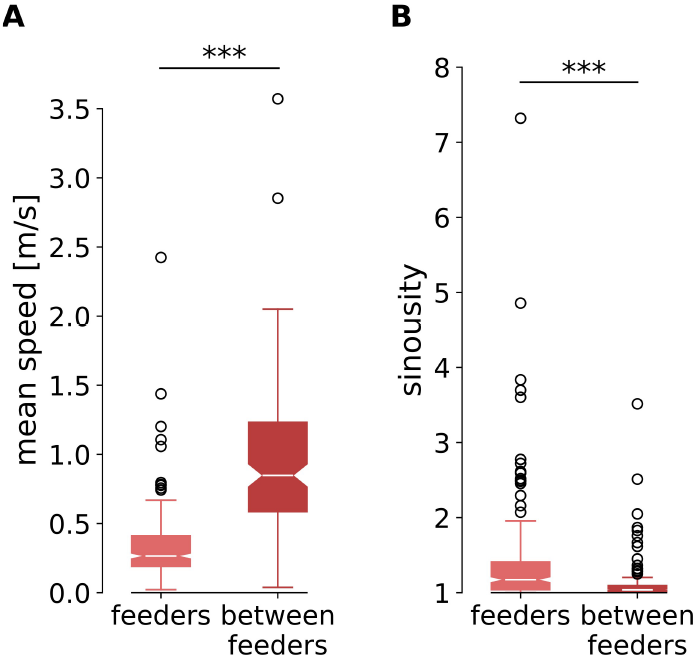
Sinuosity of the bees trajectories in **A** (1 = straight path, 1 *<* very sinuous path) and the mean speed in **B** of the bees in the areas around the feeders (light red) and between the feeders (dark red). The sinuosity is higher (Mann-Whitney *U*-test, *n*_1_ = *n*_2_ = 31, p *<* 0.001) and the speed lower in the areas around the feeders than between the feeders (Mann-Whitney *U*-test, *n*_1_ = *n*_2_ = 31, p *<* 0.001).

The search distributions of the bees around the feeder positions and the local landmark cue provided initial insights into the bees’ motivation and learning (see Fig. 8 for xz axes, see Supplementary Material for xy and yz axes). Kernel density estimation (KDE) of the bees’ search patterns showed that in the high-sphere test, the bees predominantly searched around the lowest feeder position (Fig. 8A). This suggests that they associated the feeder with the training height. However, some search activity was also observed around the intermediate and highest feeder positions, indicating that the bees did not fully discriminate between these positions based on the height. These results highlight that while the bees appeared to use relative height information between the presented sphere and the feeder to some extent, their ability to differentiate between heights was not entirely precise.

**Fig. 8.**
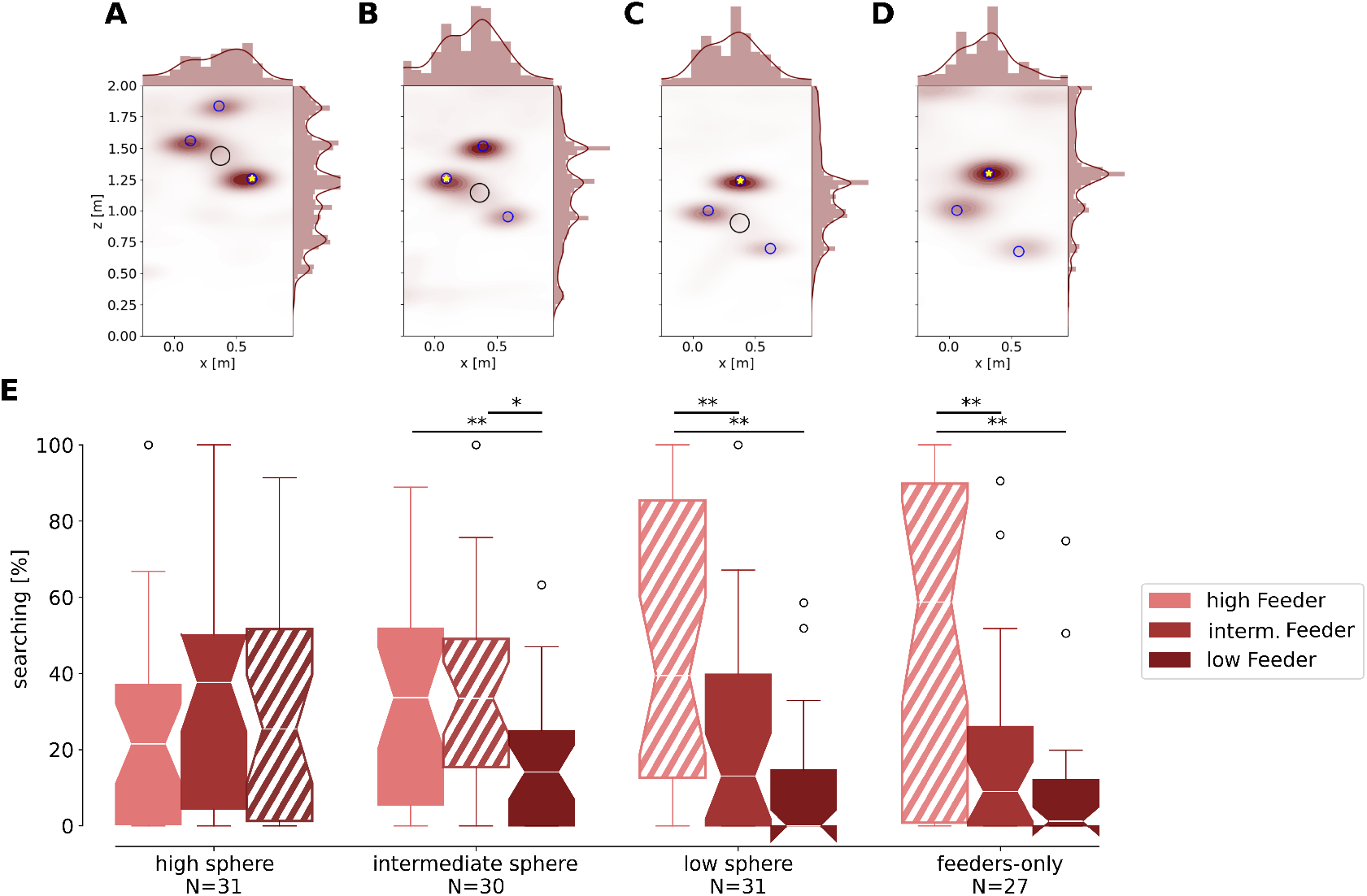
Kernel density estimations of the bees search and the comparisons of time they spent at the feeder positions in the height estimation experiment. **A-D**: Kernel density estimations of the bees search in the tests high sphere (**A**, N = 31), intermediate sphere (**B**, N = 30), low sphere (**C**, N = 31) and the test with only feeders without the low sphere (**D**, N = 27). The sphere’s position is given with a black circle, and the positions of the feeder with blue circles. The correct feeder, in respect to its height for the respective test, is indicated by a yellow star symbol. Heat maps show the highest density of bee position in red, and in white, if no bees were detected, they are seen from the x and z axes given in meters. At the top and right margins, histograms and probability density functions along the x and z axes are shown. **E**: Search proportion at the three feeders (indicated by the color rose for high, red for intermediate feeder, and dark red for the low feeder) in all four tests (high sphere, intermediate sphere, low sphere, feeders-only). The hatched box indicates the feeder at the training height. For the high sphere test, the bees (N = 31) searched a similar amount of time at the three feeders and showed a trend to search more at the lowest feeder than at the highest one. For the intermediate-sphere test, the bees (N = 30) searched more at the high and intermediate feeders than at the low feeders. For the low sphere test, the bees (N = 31) searched the most at the high feeder, which was at the training height. A similar behavior was observed for the test with only feeders without the low sphere (N = 27).

Similarly, in the intermediate sphere test, the intermediate sphere was placed in the arena center, with the feeders were placed higher or lower than the sphere or at a similar height (the training height). The KDE distributions show much search around the highest feeder and much less around the intermediate and lowest feeders (Fig. 8B). Since the intermediate feeder was positioned at the training height, the bees searched most at the feeder higher than the training height.

As we tested the high and intermediate spheres, we tested the lowest sphere. The lowest sphere was hanging in the center, and the highest feeder was placed at the training height, while the intermediate and lowest feeders were generally lower than the training height. The bees searched most at the highest feeder at training height (Fig. 8C).

To test if the bees used the sphere as a reference, we conducted a “feeders-only” test with the same three landing platforms, but no sphere. The search distributions are similar to the low-sphere test, with the bees searching at the training height (Fig. 8C&D).

All in all, the search distributions show that the bees searched for the feeder at its training height. A comparison of time spent at the three feeder heights in the four tests confirms these observations (Fig. 8E). In the high sphere test, the bees searched least at the highest feeder and a similar portion of their time at the intermediate and lowest feeder. However, no significant difference was found (ANOVA: df = 2.0, F = 0.089, Tukey: p = 0.9). In the intermediate sphere test, bees spent more time searching at the high and intermediate feeders than at the low feeder, but no significant difference was found between the high and intermediate feeders (ANOVA: df = 2.0, F = 5.798, Tukey: p _high-low_ = 0.021, p _intermediate-low_ = 0.007). In the low sphere test, we found a significant difference between the time spent at the highest feeder and at the lowest feeder, indicating that the bees could discriminate between the highest feeder, at the training height, and the lowest feeder (ANOVA: df = 2.0, F = 14.287, Tukey: p = _high-intermediate_ = 0.003, p _high-low_ = 0.001). The search proportion at the intermediate feeders shows a trend of less search than at the highest feeder. In the “feeders-only” test, the bees spent most time searching at the highest feeder and significantly less at the other two feeders (ANOVA: df = 2, F = 15.398, Tukey: p _high-intermediate_ = 0.001, p _high-low_ = 0.001).

Since the bees chose most clearly the correct height when the constellation of feeders without the sphere being placed closest to the ground floor, we assume that the bees used the ground floor to estimate their absolute height, most likely based on ventral optic flow. The single spheres that were placed higher (high and intermediate tests) seemed to act like distractors for height estimation. Additionally, the absolute height estimation based on ventral optic flow might not have worked precisely due to the larger distance between the ground floor and the landing platforms. Taken together, we found that bees can find a food location only based on its height, and they would use absolute distance to the ground rather than their spatial relation to the landmarks.

## DISCUSSION

We investigated how bees locate a goal in 3D space when the goal location is indicated by surrounding local landmark cues, i.e. spheres placed at different locations in 3D space around a feeder. We found that the bees associated the cues with the goal, similar to 2D paradigms. However, we did not observe a concentrated search around the feeder position as found in those studies investigating the goal localization of a food goal in 2D (Cartwright et al., 1983; Cheng et al., 1987). In the 2D experiments, when the feeder was removed, the bees searched around this position, but in our 3D, the bees did not search precisely at the location where the feeder was positioned in the training situation. This suggests that bees do not - as is generally assumed for the corresponding 2D situation - use the positions of the landmarks alone to localize a goal position, if the goal itself is not visible. Since the animals still coarsely search for the goal location with reference to the landmarks, we hypothesize that they might need additional visual cues to precisely determine the search area.

To explore what additional cues the bees might use to choose a search area, we further assessed the ability of bumblebees to learn the height of a food source. We found that bees could locate the goal’s height using global cues like the ground floor, likely relying on ventral optic flow to estimate distance to the ground. However, the local landmarks and greater distance from global cues acted as distractors, reducing the accuracy of their search.

### Differences in 2D and 3D

Transferring an experimental paradigm from 2D to 3D is not trivial. We found that the bees did not exhibit such a clear search pattern in 3D as shown in 2D (Cheng et al., 1987). A major difference is that the bees can land in the 2D paradigms even if they cannot find their target on the ground, whereas this was not possible in our corresponding experiment in 3D. Therefore, in the second experiment, we provided the bees with alternative landing platforms at different positions relative to the landmarks. This showed that the height of the goal is an important source of information, in addition to the landmark information. This issue is crucial for understanding how bees navigate in 3D, as 3D environments add more challenges than 2D. Bees engage in various flight maneuvers, including pivoting, turning, sideways movement, hovering, and backward flight (e.g. Linander et al., 2018; Doussot et al., 2021). This dynamic range of movements suggests that bees adjust their body and head positions to memorize views that guide them back to nest locations or other goals (Doussot et al., 2021). Previous studies have highlighted the importance of this dynamic behavior, indicating that bees employ spatial awareness strategies to navigate effectively in complex 3D landscapes (Linander et al., 2018). Our findings further contribute to this understanding, revealing that while bees demonstrate the ability to localize goals vertically, success is limited when the landmark-goal arrangement is closer to ground level. This insight emphasizes the nuanced challenges bees face in 3D navigation and the importance of considering spatial context in understanding their behaviors.

### Use of optic flow for flight control

In the height estimation experiment, we observed that the bees searched around the trained feeder height without requiring the use of local landmarks. This suggests that the bees used ventral optic flow for goal height estimation. The utilization of optic flow for flight control is a crucial aspect of bees’ navigation in 3D (Lecoeur et al., 2019; Egelhaaf, 2023). Extensive studies have demonstrated that flying insects rely heavily on optic flow to control their flight (Linander et al., 2018; Frasnelli et al., 2021). Bumblebees, in particular, prioritize motion cues from the ground, using ventral optic flow to control their altitude and lateral position (Linander et al., 2017). When the availability of ventral optic flow cues is limited, bumblebees adjust their flight height to maintain the ability to resolve ground texture (Portelli et al., 2010; Linander et al., 2018). Since the bees required a relatively close distance between the goal and the ground (up to 1.25 m) to accurately estimate goal height in our experimental setup, we assume they relied on ground-based allocentric distance estimation using optic flow. Srinivasan et al. (1989) demonstrated that honeybees can distinguish height differences between objects of more than 2 cm based on optic flow, while our experiment suggests an upper limit of 1.25 m for ground-based distance estimation.

The upper limit of 1.25 m for bumblebees’ distance estimation in our study can likely be explained by the relationship between optic flow and altitude. At a given flight speed, optic flow decreases with height, and due to inevitable signal-to-noise limitations in neural processing, it must remain above a certain threshold to be detectable. Electrophysiological studies in blowflies have examined this in detail, showing that at a flight speed of approximately 0.5 m/s, distances of up to 1 m can be reliably determined based on optic flow (Kern et al. 2005). Thus, height estimation using optic flow is only feasible if the animal flies sufficiently close to the ground. Given that the flight speeds of bees during search at and between the feeder (Fig. 7A) are comparable to those observed in blowfly experiments (Kern et al., 2005) - and that the spatial resolution of the bee’s eye is similar to that of blowflies (Petrowitz et al., 2000; Spaethe et al., 2003) - the most plausible explanation for the observed height limit in our experiments is the reliability of optic flow detection and neural processing.

### Local and global cues

Our results on the search behaviour of bees for goal locations indicate that bumblebees rely on both local and global visual cues for navigation. Cheng et al. (1987) compared the use of global and more local cues. They placed cylindrical landmarks at various distances to the goal and observed the honeybees’ search patterns. The bees were found to search more at the landmarks closer to the goal, even when these were moved in relation to the more global landmarks. Transferred to our 3D-setting, one would have expected that the local landmark cues, i.e. spheres placed in 3D, should have played a more significant role than more global reference structures of the arena, such as the ceiling or optic flow induced by the ground. The use of cues for goal localization on the horizontal plane appears to differ from that along the vertical axis. Additionally, a constant reference like the ground might be more reliable than more local landmarks, pointing to a similar strategy as shown a higher weighting of stationary objects by humans (Roy et al., 2022).

## Conclusion

When challenged during foraging in a 3D world where the height of a food source and landmark cues also play a role, bumblebees can learn and return to a specific flower height. While bees primarily searched near the landmark constellation, they tended to undershoot the trained goal position. Their height estimation was more accurate when the food source and landmarks were closer to the floor, suggesting they use optic flow to estimate goal’s height. Further studies should investigate how the bees sample optic flow for height estimation, for example, by systematically and periodically changing their flight altitude (Bergantin et al., 2021). Additionally, it would be interesting to know which information the bees would need to locate the correct height when the feeder and landmarks are farther away from the ground floor. Manipulating the optic flow by varying patterns, indicating distances different from the learnt one, is needed to strengthen our hypothesis.

## Acknowledgements

We would like to thank Leo Werner, Paula Bräuer und Madelene Dombrowski for their help during the data collection, and Charlotte Doussot for designing the flight arena and fruitful discussion during the study.

## Contribution

AS: conceptualisation, investigation, data curation, formal analysis, methodology, visualisation, writing - original draft, review, and editing. OJNB: conceptualisation, software, supervision, project administration, writing - review, and editing. ME, ML: funding acquisition, project administration, resources, supervision, writing - review, and editing

## Data availability

The data-sets and analysis pipeline for this study can be found in the repository “3D goal localization in bumblebees”.

