## Supplementary Material for "Bumblebees locate goals in 3D with absolute height estimation from ventral optic flow"

### 1 SUPPLEMENTARY FIGURES

#### 1.1 Angular object size

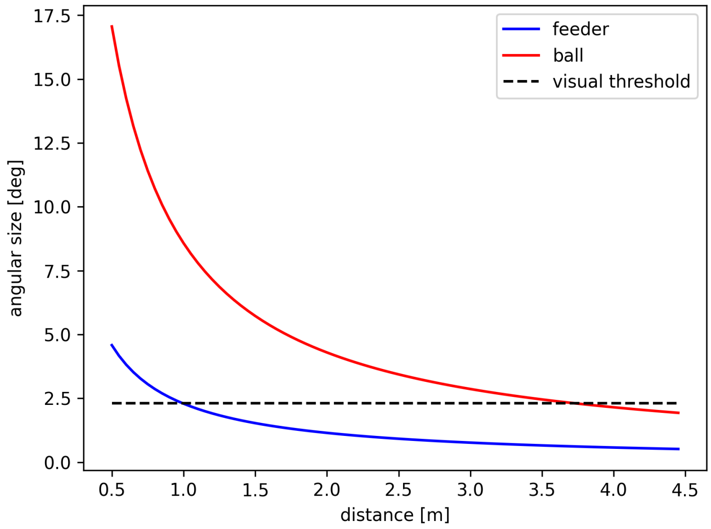


**Figure S1.** Angular size of the objects (spheres, red line, and feeder, blue line) in relation to the distance between the object and the observer. The visual threshold of 2.3 degrees for object detection in bumblebees is indicated by the black dashed line.

#### 1.2 Photographs of the setup

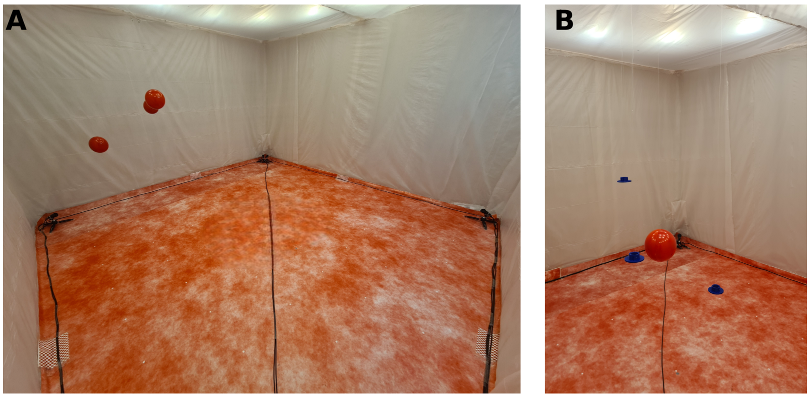


**Figure S2. Photographs of the experimental setups as seen from the bee's nest entrance. A**: Photograph of the experimental setup of the 3D goal localisation experiment shows the three sphere at the training position with the feeder being removed (control test). **B**: Photograph of the experimental setup of the height estimation experiment shows the condition of the low sphere test with the low sphere surrounded by three feeders.

### 1.3 3D goal localization

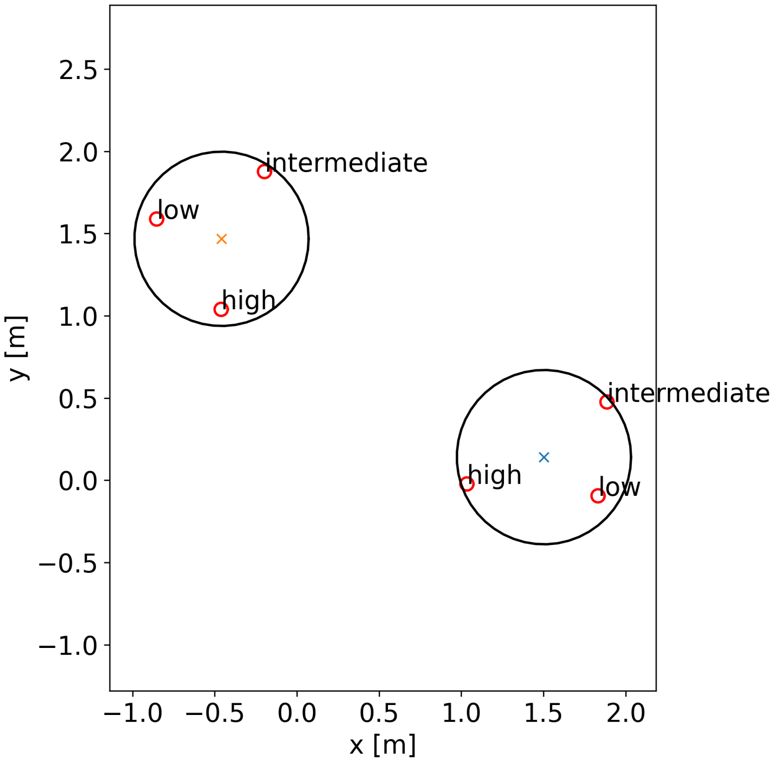


**Figure S3. Schematic showing the area used for comparing the time the bees spent around the position of the feeder during training and during test.** The schematic shows the area with a radius of 0.53m around the position of the feeder during training and during test that was used to compare the time the bees spent at these two locations during the test. The spheres are indicated by red circles and the position of the two feeders are respectively indicated by a cross (yellow for the training position and blue for the test position).

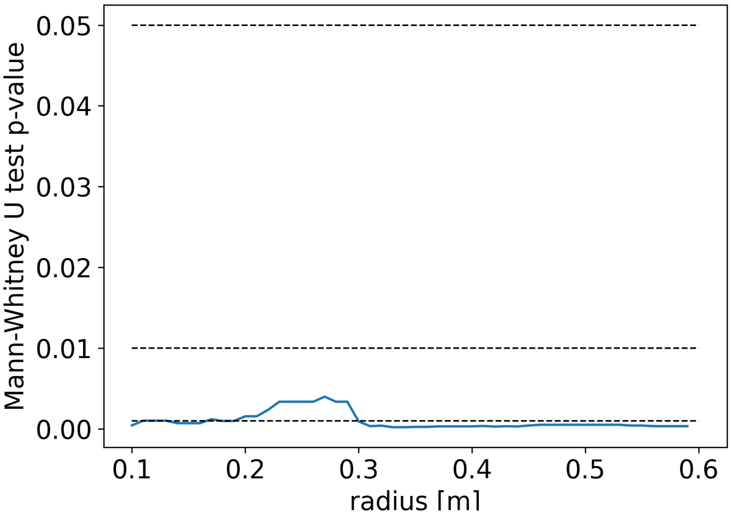


**Figure S4. Mann-Whitney U test with varying radii around the feeder position during training and during test of the bees spent at these locations.** Mann-Whitney U test p-values (blue line) were calculated for varying radii of the volume around the feeders' positions (0.1 to 0.6 m) to analyze the time spent at the training and at the test location of the feeders. The reference to the significant levels (0.05, 0.01, 0.001) are shown as black dashed lines. The time the bees spent at the test position of the feeder is for all radii significantly different from the time spent at the training position.

### 1.4 Height estimation

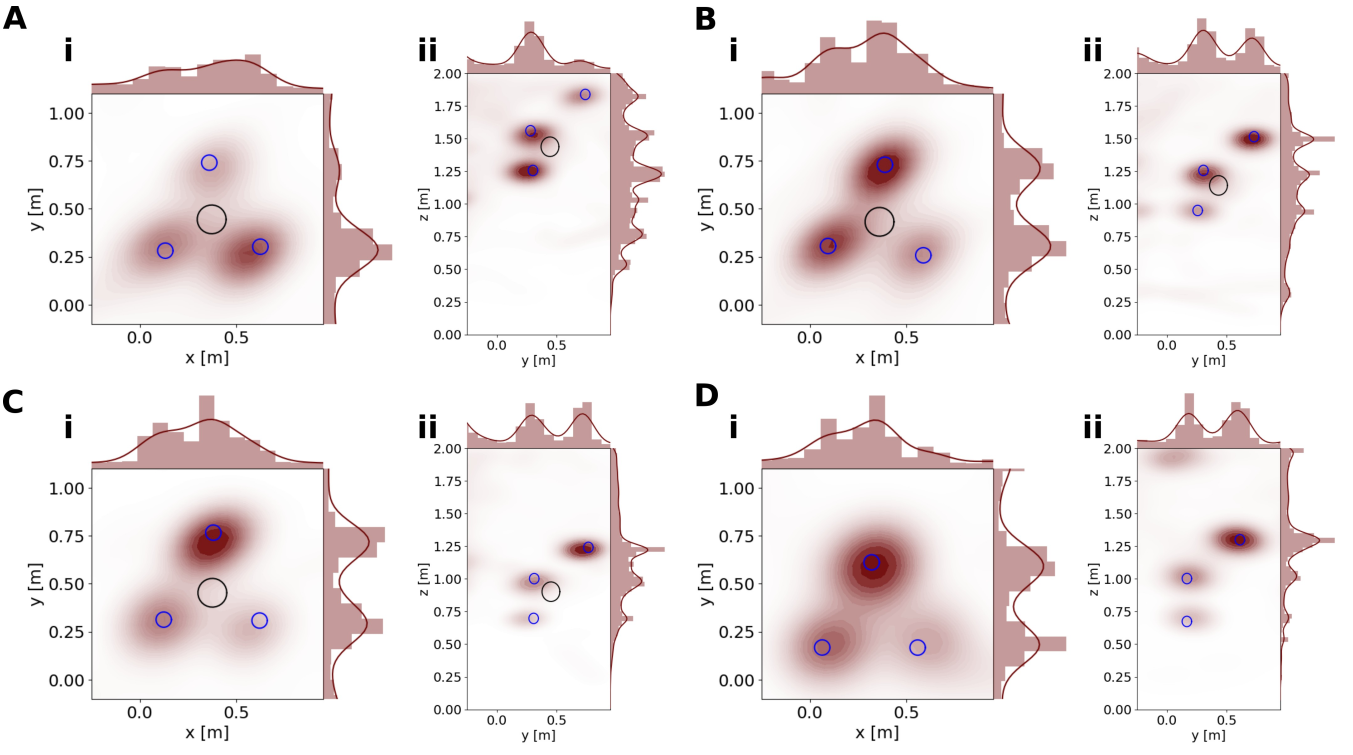


**Figure S5.** Kernel density estimation of the search distributions in the height estimation experiment. The tests high sphere (**A**), intermediate sphere (**B**) low sphere (**C**) and feeders-only (**D**) are presented along the axes x-y (**i**) and y-z (**ii**). The sphere's position is given with a black circle, and the positions of the feeder with blue circles. The correct feeder, in respect to its height for the respective test, is indicated by a yellow star symbol. Heat maps show the highest density of bee position in red, and in white, if no bees were detected, they are seen from the x and y axes in **i** and y and z axes in **ii** given in meters. At the top and right margins, histograms and probability density functions along axes are shown.
